# Deciphering the rules underlying xenogeneic silencing and counter-silencing of Lsr2-like proteins

**DOI:** 10.1101/751156

**Authors:** Johanna Wiechert, Andrei Filipchyk, Max Hünnefeld, Cornelia Gätgens, Ralf Heermann, Julia Frunzke

## Abstract

Lsr2-like nucleoid-associated proteins play an important role as xenogeneic silencers (XS) of horizontally acquired genomic regions in actinobacteria. In this study, we systematically analyzed the in vivo constraints underlying silencing and counter-silencing of the Lsr2-like protein CgpS in *Corynebacterium glutamicum*. Genome-wide analysis revealed binding of CgpS to regions featuring a distinct drop in GC-profile close to the transcription start site (TSS), but also identified an overrepresented motif with multiple A/T steps at the nucleation site of the nucleoprotein complex. Binding of specific transcription factors (TFs) may oppose XS activity leading to counter-silencing. Following a synthetic counter-silencing approach, target gene activation was realized by inserting operator sites of an effector-responsive TF within various CgpS target promoters resulting in an increased promoter activity upon TF binding. Analysis of reporter constructs revealed maximal counter-silencing when the TF operator site was inserted at the position of maximal CgpS coverage. This principle was implemented in a synthetic toggle switch, which features a robust and reversible response to effector availability highlighting the potential for biotechnological applications. Altogether, our results provide comprehensive insights into how Lsr2 silencing and counter-silencing shapes evolutionary network expansion in this medically- and biotechnologically-relevant bacterial phylum.

**IMPORTANCE:** In actinobacteria, Lsr2-like nucleoid-associated proteins function as xenogeneic silencers (XS) of horizontally acquired genomic regions, including viral elements, virulence gene clusters in *Mycobacterium tuberculosis*, and genes involved in cryptic specialized metabolism in *Streptomyces* species. Consequently, a detailed mechanistic understanding of Lsr2 binding in vivo is relevant as a potential drug target and for the identification novel bioactive compounds. Here, we followed an in vivo approach to investigate the rules underlying xenogeneic silencing and counter-silencing of the Lsr2-like XS CgpS from *Corynebacterium glutamicum*. Our results demonstrated that CgpS distinguishes between self and foreign by recognizing a distinct drop in GC-profile in combination with a short, sequence specific motif at the nucleation site. Following a synthetic counter-silencer approach, we studied the potential and constraints of transcription factors to counteract CgpS silencing thereby facilitating the integration of new genetic traits into host regulatory networks.

## INTRODUCTION

Horizontal gene transfer (HGT) is a major driver of bacterial evolution and plays an important role in creating genetic diversity (1). The rapid acquisition of new beneficial traits can create a competitive advantage for the recipient cells (1, 2). However, the chance that foreign DNA decreases the fitness of the cell is high, since it may lead to interference with regulatory networks, high transcriptional and translational costs, sequestration of cellular machineries and cytotoxic gene products (3–8). Therefore, bacteria evolved a variety of immune systems allowing them to deal with foreign DNA (9). CRISPR-Cas and restriction modification systems are nuclease-based defense mechanisms enabling the recognition and targeted degradation of invading DNA (10–12). In contrast to these destructive immune systems, xenogeneic silencing enables the tolerance of foreign DNA and consequently fosters the acquisition of novel genetic material into the host chromosome (13). Xenogeneic silencing is based on specific nucleoid-associated proteins (NAPs), so-called xenogeneic silencers (XS) (3). Known XS proteins belong to one of four currently described classes: H-NS-like proteins of proteobacteria like *Escherichia coli*, *Yersinia* and *Salmonella* (14–16), MvaT/U-like proteins found in gamma-proteobacteria of the Pseudomonodales order (17), Lsr2-like XS of Actinomycetes (18, 19) and Rok present in different bacilli, including *Bacillus subtilis* (20, 21). Although XS were convergently evolved and show only low sequence similarity, the domain properties of their N-terminal oligomerization domains and their C-terminal DNA-binding domains are similar (19, 20, 22, 23). Their binding mechanisms are diverse but they all preferentially bind to horizontally acquired DNA, which has typically a higher AT-content than the genome of the recipient cell (4, 24). The broad distribution of XS among prokaryotes emphasizes the strong need to discriminate between “self” and “non-self” across phylogenetic clades (13). Even so the GC-content of microbial genomes dramatically varies from 75% (Actinobacteria) to less than 20% (bacterial endosymbionts) (25, 26), horizontally acquired regions typically feature a lower GC-content than their resident genome emphasizing base composition as a major discrimination factor shaping microbial genome evolution (3).

Several studies based on variants defective in oligomer formation revealed that binding of XS proteins to the DNA alone is insufficient for silencing (27–29). The formation of higher order nucleoprotein complexes, instead, mediates silencing of the target genes by occlusion or trapping of the RNA-polymerase, by interference with the transcription elongation complex or by enhancing termination (30, 31). To get access to potentially encoded beneficial traits, cells must integrate foreign genes into pre-existing regulatory circuits allowing their controlled expression at appropriate time points and physiological or environmental conditions (32, 33). In contrast to classical activation, counter-silencing is based on the interference of a DNA-binding protein, e.g. a transcription factor (TF), with the silencer-DNA complex leading to transcription initiation without depending on the direct interaction with the RNA-polymerase (32, 33). Counter-silencing of H-NS was addressed by several studies either by following a synthetic approach at well-studied promoters (34, 35) or by the analysis of the promoter architectures in the PhoPQ regulatory network (33). This recent study by Will et al. emphasizes that the principle of H-NS xenogeneic silencing and counter-silencing provides a certain degree of flexibility fostering evolutionary network expansion (33).

Compared to H-NS in proteobacteria, much less is known about Lsr2-like XS proteins conserved throughout the actinobacteria. In *Mycobacterium tuberculosis*, Lsr2 acts as a master regulator of multiple virulence-associated genes (19, 22) and was suggested to be involved in the manifestation of multi-drug tolerance (36). The essentiality of Lsr2 for this human pathogen makes this XS protein a highly promising drug candidate (37). In *Corynebacterium glutamicum* the Lsr2-like XS protein CgpS was also shown to play an essential role as a silencer of cryptic prophage elements, whose entrance into the lytic cycle would otherwise cause cell death (4, 38). In contrast to mycobacteria and corynebacterial species, *Streptomyces* typically encode two Lsr2-like proteins. Here, the prototypical *lsr2* gene, showing the highest sequence identity to mycobacterial Lsr2, was recently described to silence the expression of specialized metabolic clusters (39). Considering the important role of Lsr2 proteins in the medically and biotechnologically important phylum of actinobacteria, a detailed mechanistic understanding of Lsr2 binding in vivo is relevant as a potential drug target and for the identification novel bioactive compounds.

In this study, we set out to systematically assess the rules underlying silencing and counter-silencing of Lsr2-like XS by using the Lsr2-like protein CgpS of *Corynebacterium glutamicum* as a model (4). To the best of our knowledge, it is the first detailed analysis of the counter-silencing mechanism of an Lsr2-like XS protein. Bioinformatic analysis of CgpS ChAP-seq (chromatin affinity purification and sequencing) data revealed a clear preference of CgpS towards AT-rich stretches containing A/T steps (alternation of A to T and vice versa). In vivo reporter studies with synthetic promoter variants verified the importance of a distinct drop in GC-profile and revealed the overrepresentation of a short, sequence specific motif at CgpS target regions. Insertion of TF operator sites at different positions within various CgpS target promoters was shown to counteract CgpS silencing, showing the most prominent effect at the position of maximal CgpS binding. With this approach, we provide important insights into the in vivo constraints of Lsr2 counter-silencing and contribute to an understanding of how bacteria can evolve control over the expression of horizontally acquired genes.

## RESULTS

### In vivo analysis of CgpS binding preferences

Recent genome-wide profiling studies revealed that the Lsr2-like xenogeneic silencer CgpS preferentially binds to AT-rich DNA sequences in the genome of *C. glutamicum* ATCC 13032 (4). To determine the parameters affecting CgpS binding and silencing in vivo, we systematically analyzed the peak sequences obtained from CgpS ChAP-seq analysis (4) and subsequently confirmed our conclusion by testing the silencing of synthetic promoter variants. Remarkably, an overlay of the GC-profiles of all 35 CgpS target promoters located within the prophage element CGP3 revealed a high degree of similarity with a distinct drop in GC-content matching the position of maximal CgpS coverage (Figure 1A). Genome-wide analysis of AT-rich genomic regions revealed that the fraction of CgpS-bound sequences increased with the length of the particular AT-stretch. While increasing numbers of G/C interruptions negatively influenced the proportion of CgpS-bound targets (Figure 1B), a higher number of A/T steps (alternation of A to T and vice versa) increased the fraction of CgpS-bound sequences by trend (Figure 1C). This trend became especially evident in the case of AT-rich stretches of medium length (14-30 bp).

**Figure 1:**
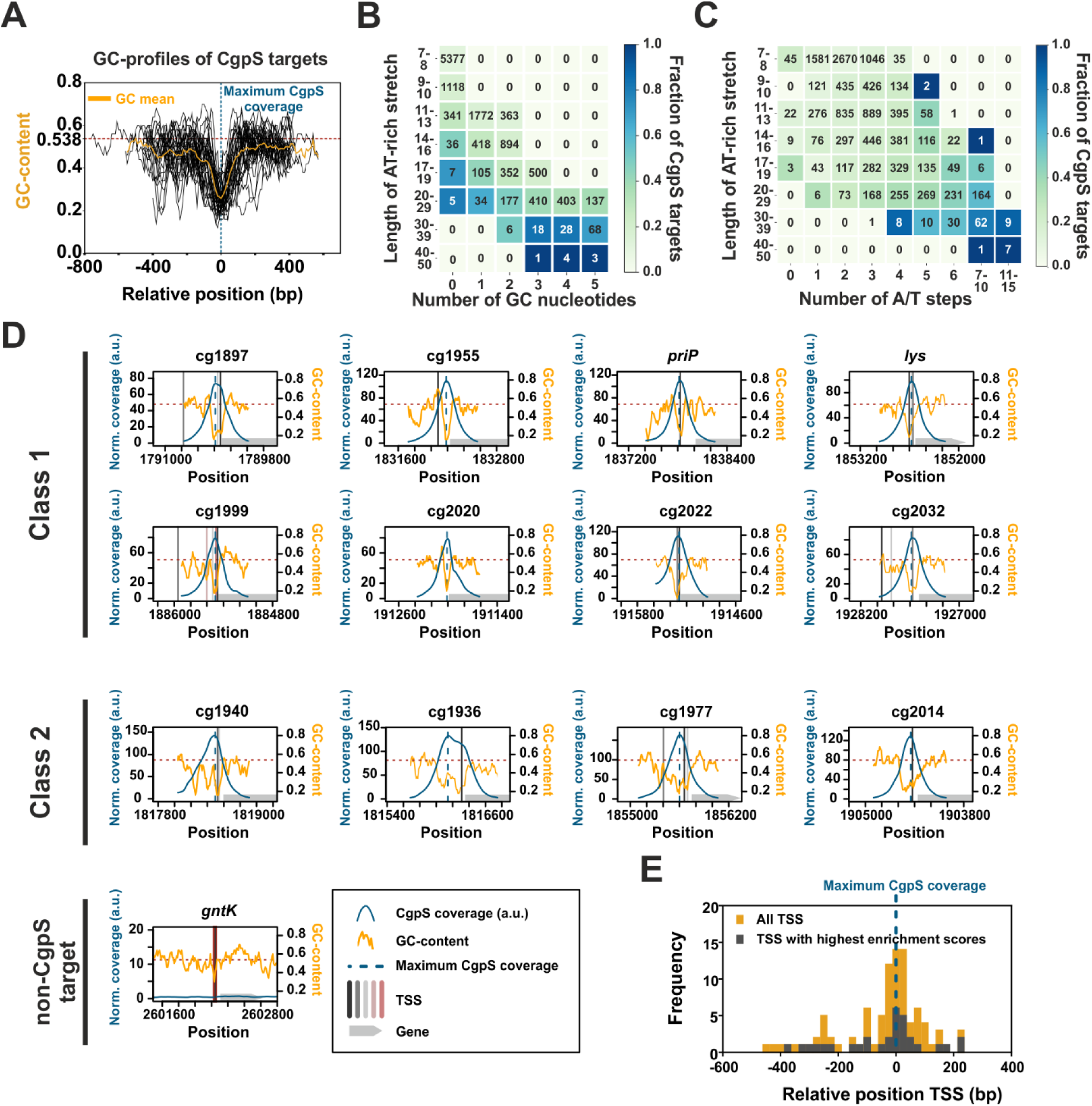
CgpS preferentially binds to long and consecutive AT-stretches. **A)** Overlay and calculated mean (orange curve) of GC-profiles of CgpS target promoters located within the CGP3 prophage (n=35) (4). Profiles were calculated by a rolling mean with a window size of 50 bp and a step size of 10 bp. The GC-profiles of the promoters were normalized regarding the orientation and position of the maximal CgpS binding peak (blue line), which was defined for all sequences as position 0. The mean GC-content of the *C. glutamicum* genome (70) is shown as red line (53.8%). **B/C)** Genome-wide analysis of CgpS binding to consecutive AT-stretches of different lengths considering G/C interruptions (B) or number of A/T steps (allowing up to 5 G/C interruptions). A/T steps are defined as alternations of A to T and vice versa (C). The value in the array represents the number of stretches found in the *C. glutamicum* genome fitting the respective criteria, while the color indicates the fraction of CgpS targets per array. **D)** Divergent correlation of GC-profiles and CgpS coverage of CgpS target promoters. CgpS coverage obtained from previous ChAP-seq experiments (4) was calculated with a rolling mean with a window size of 50 and a step size of 10. All identified TSS (see material and methods and the supplemental methods) are shown in Table S1 and represented as vertical black, grey and red lines (mapped according to their enrichment scores: black > shades of grey > red). Positions of maximal CgpS coverage and average GC-content are shown as described in (A). The corresponding genes are shown as grey arrows. Promoters were grouped into two classes based on the width of the region bound by CgpS (class 1 promoters: 500-850 bp, typically featuring one distinct drop in GC-profile; class 2 promoters: >850 bp, often broader and containing multiple drops in GC-content). As a negative control, the non-CgpS target promoter of the gene *gntK* is shown. **E)** Frequency distribution of relative positions of all new identified TSS (yellow) of CgpS target promoters referred to the position of maximal CgpS binding. TSS showing the highest enrichment scores per gene are highlighted in grey.

Overall, this analysis suggested that long and consecutive AT-stretches represent the main determinant of CgpS target binding. Individual inspection of CgpS targeted phage promoters revealed a significant correlation between the CgpS peak maximum and the GC minimum in this area (Figure 1D). Depending on the widths of the CgpS coverage peaks, promoters were grouped into two classes. Class 1 consists of promoters with peak widths between 500 and 850 bp which typically show one distinct drop in GC-profile, while CgpS coverage peaks of class 2 promoters are wider than 850 bp and the corresponding GC profiles often feature broader or multiple drops.

Due to efficient CgpS-mediated silencing of gene expression, most transcriptional start sites (TSS) of CgpS target promoters had not been identified in previous studies (40). It represents, however, an advantage of the chosen model system that expression of the majority of CgpS targets can be induced by triggering prophage induction using the DNA damaging antibiotic mitomycin C. To provide comprehensive insights into the promoter architecture of CgpS targets, TSSs were determined under conditions triggering phage gene expression (600 nM mitomycin C). Strikingly, this analysis revealed that in the majority of CgpS target promoters TSSs are located close to the position of maximal CgpS coverage and GC minimum (Figure 1D, E, see Table S1 for the complete dataset).

### Design, build & test - Relevance of a DNA motif for CgpS binding and silencing

Bioinformatic analysis of CgpS target sequences confirmed the preference of CgpS for AT-rich DNA-sequences. However, neither the distinct drop in GC-content nor the occurrence of long and consecutive AT-stretches were unique to CgpS targets (Figure 1) indicating that additional parameters support CgpS to specifically recognize its targets. Interestingly, a MEME-ChIP analysis (41) on CgpS bound DNA sequences revealed a 10 nucleotides long AT-rich binding motif (E-value: 5.2 * 10^-9^) containing A/T steps (Figure 2A), which was found in 51 of 54 bound promoter regions. Remarkably, the presence of this motif within AT-rich stretches of different lengths significantly increased the fraction of CgpS bound sequences by a factor of up to 2.8-fold (Figure 2A). However, the genome-wide search for motif occurrence using the online tool FIMO (*Find Individual Motif Occurrences*) (42) revealed that about 85% of the motifs (669/785) within the *C. glutamicum* genome were not bound by CgpS indicating that the motif alone is not sufficient to permit CgpS binding.

**Figure 2:**
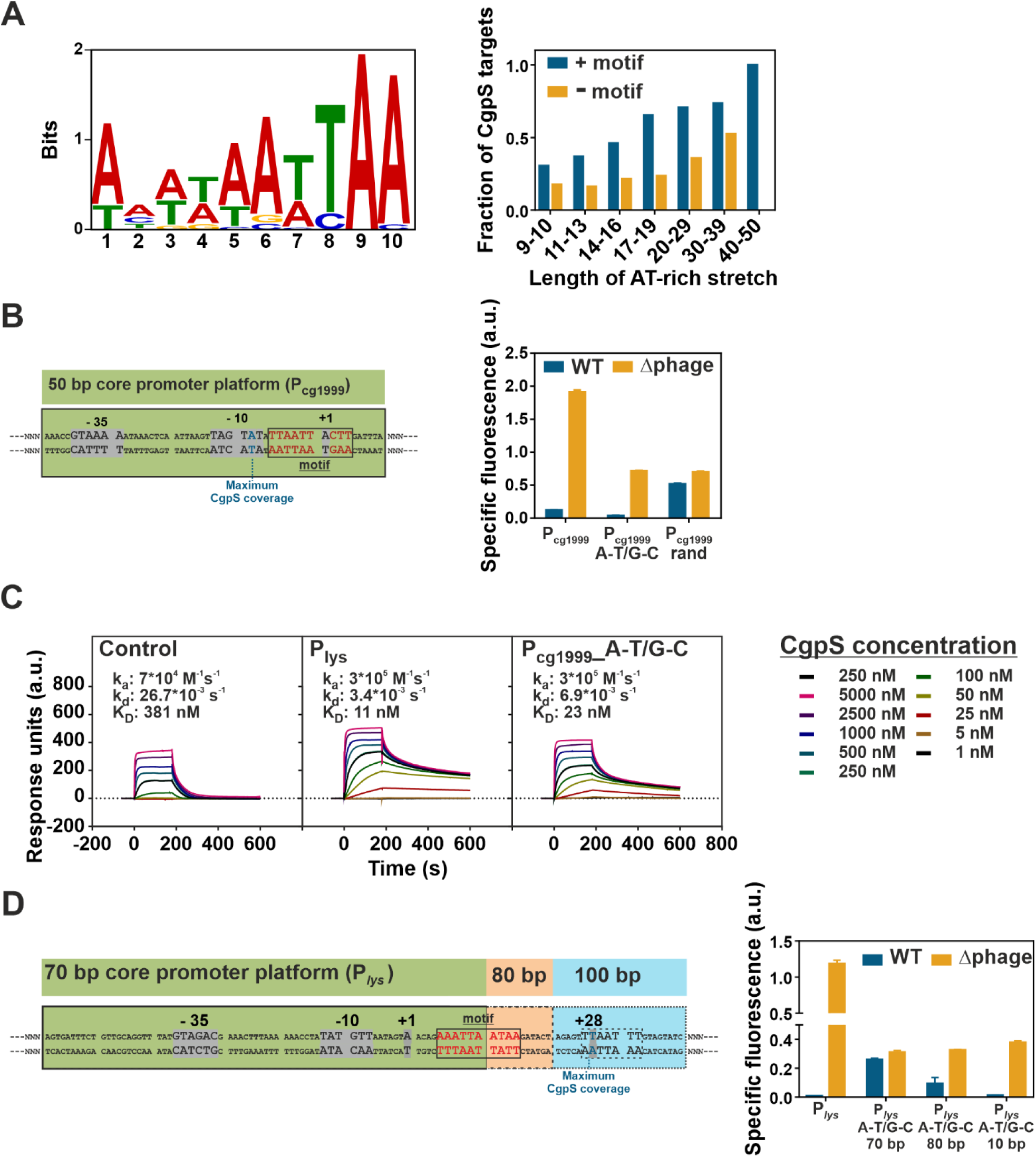
A synthetic in vivo approach to dissect the relevance of the GC-profile and a sequence-specific binding motif for CgpS silencing. **A)** Identified 10 bp CgpS binding motif using MEME-ChIP (41) analysis found within 51 of 54 CgpS target promoters (4) (E-value: 5.2 * 10^-9^). Bar plot represents the genome-wide fraction of CgpS targets in AT-stretches of different length allowing up to 5 G/C interruptions with or without the identified motif. **B)** CgpS silencing of synthetic constructs (P_cg1999__A-T/G-C and P_cg1999__rand) based on a 50 bp core promoter region (green box) of the phage gene cg1999. The fixed 50 bp DNA sequence covered the -10 and -35 box, the positions of TSS (40) and potential binding motif (grey box). The adjacent sequence (N upstream: 260 bp, N downstream 48 bp) was either adjusted to maintain the native density of AT-stretches (P_cg1999__A-T/G-C: exchange of A to T and G to C) or was randomized (P_cg1999__rand). **C)** Surface plasmon resonance analysis of CgpS binding to the synthetic promoter P_cg1999__A-T/G-C (423 bp) in comparison to the negative control P_cg3336_ (424 bp) and the native CgpS target P*_lys_* (424 bp). **D)** CgpS silencing of synthetic constructs based on fixed 70-100 bp promoter regions of the phage gene *lys*. The 70 bp sequence (green box) covered the -10 and -35 box, TSS but only half of the putative motif. The 80 bp region (green and orange boxes) covered the motif completely and the 100 bp region (all boxes) additionally covered the position of maximal CgpS coverage. The adjacent sequences (N upstream: 304 bp, N downstream 70-100 bp) were adjusted to maintain the native density of AT-stretches (A-T/G-C). **B/C)** Shown are reporter outputs (Venus) of native and the corresponding synthetic variants (plasmid backbone pJC1) in wild type and Δphage strain (Δ*cgpS*) after 5 hours of cultivation in a microtiter cultivation system in CGXII medium containing 100 mM glucose. Shown are mean values and standard deviation of biological triplicates. All synthetic sequences are listed in Table S2H.

In the following we used an in vivo approach, to test the hypothesis whether the combination of the motif and the drop in GC-profile are sufficient for CgpS-mediated silencing of gene expression. For this purpose, different synthetic promoter variants were designed based on the 50-70 bp core promoter regions of phage genes (P_cg1999_ and P*_lys_*). In the case of P_cg1999_, the DNA sequence containing the core promoter elements (-10 and -35 box, TSS) and the predicted binding motif (shown in Figure 2A) was kept constant (Figure 2B). The adjacent sequence was either designed to mimic the native GC-profile of P_cg1999_ (exchange of A to T and G to C and vice versa: P_cg1999__A-T/G-C) or contained a randomized sequence varying in GC-profile and sequence (P_cg1999__rand). The resulting promoter designs were fused to a gene encoding the yellow-fluorescent protein Venus. In line with our hypothesis, the construct P_cg1999__A-T/G-C featuring the native GC-profile was efficiently silenced by CgpS in the wild type strain and displayed even lower reporter output than the native phage promoter P_cg1999_ (Figure 2B). SPR analysis revealed similar CgpS binding kinetics and affinities for this synthetic promoter (K_D_=23 nM) compared to the native CgpS target promoter P*_lys_* (K_D_=11). CgpS also interacted with the control promoter fragment P_cg3336_, but with much lower affinity (381 nM) and very fast dissociation rates (Figure 2C). In the prophage-free strain Δphage, which is lacking the phage-encoded *cgpS* gene, the reporter output was significantly higher for all tested promoter fusions confirming that all designs functionally drive transcription. Silencing of the promoter variant with randomized adjacent flanks was strongly impaired demonstrating that the motif containing 50 bp core promoter region alone did not mediate silencing. This highlights the importance of the overall drop in GC-content observed at CgpS target promoters (Figure 2B).

The relevance of the identified motif was verified using synthetic promoter designs of the phage promoter P*_lys_*. Here, constructs carrying only parts of the predicted motif (70 bp core) did not permit silencing, while constructs covering the motif entirely enabled silencing (Figure 2D). In all P*_lys_*-based synthetic constructs the native GC-profile of the sequence flanking the core promoter region was mimicked but the DNA sequence was changed (A to T and G to C and vice versa). This in vivo analysis of synthetic phage promoter variants revealed that efficient CgpS silencing depended on both specific DNA sequences (binding motif) and the drop in GC-profile in the flanking regions.

### Synthetic disruptive counter-silencing

Disruptive counter-silencing was previously described as a mechanism, which may provide access to horizontally acquired genes silenced by nucleoid-associated proteins (32). To study the potential and constraints of evolutionary network expansion by counter-silencing of CgpS target promoters, a synthetic counter-silencer design was applied in this study (Figure 3A). At native target promoters (e.g. P_cg1999_ or P*_lys_*), oligomerization of the xenogeneic silencer CgpS leads to the formation of a nucleoprotein complex inhibiting transcription (4, 31). In the following, we used a set of 12 different phage promoters as a basis for synthetic counter-silencer (CS) constructs and inserted the operator sequence of an effector-responsive transcription factor (TF) into the silenced promoter regions. We postulated, that binding of the TF to its operator sequence would interfere with the silencer nucleoprotein complex and thereby mediate counter-silencing (Figure 3A). To avoid interference of the inserted operator site with CgpS-mediated silencing, we chose the operator site of the functionally redundant TFs GntR1 (Cg2783) and GntR2 (Cg1935) (summarized as GntR in the following), which bind to a well-defined short (15 bp) and AT-rich (GC-content: 27%) DNA motif (43). One of the native targets of GntR is the promoter of the *gntK* gene, which is repressed by binding of GntR. The P*_gntK_* promoter and the synthetic promoter constructs were fused via a consistent linker containing a ribosomal binding site to the reporter gene *venus* and were inserted into the plasmid pJC1. The effector molecule gluconate was shown to act as an inducer triggering the dissociation of GntR from its operator site (43), consequently leading to de-repression of P*_gntK_* (Figure 3B).

**Figure 3:**
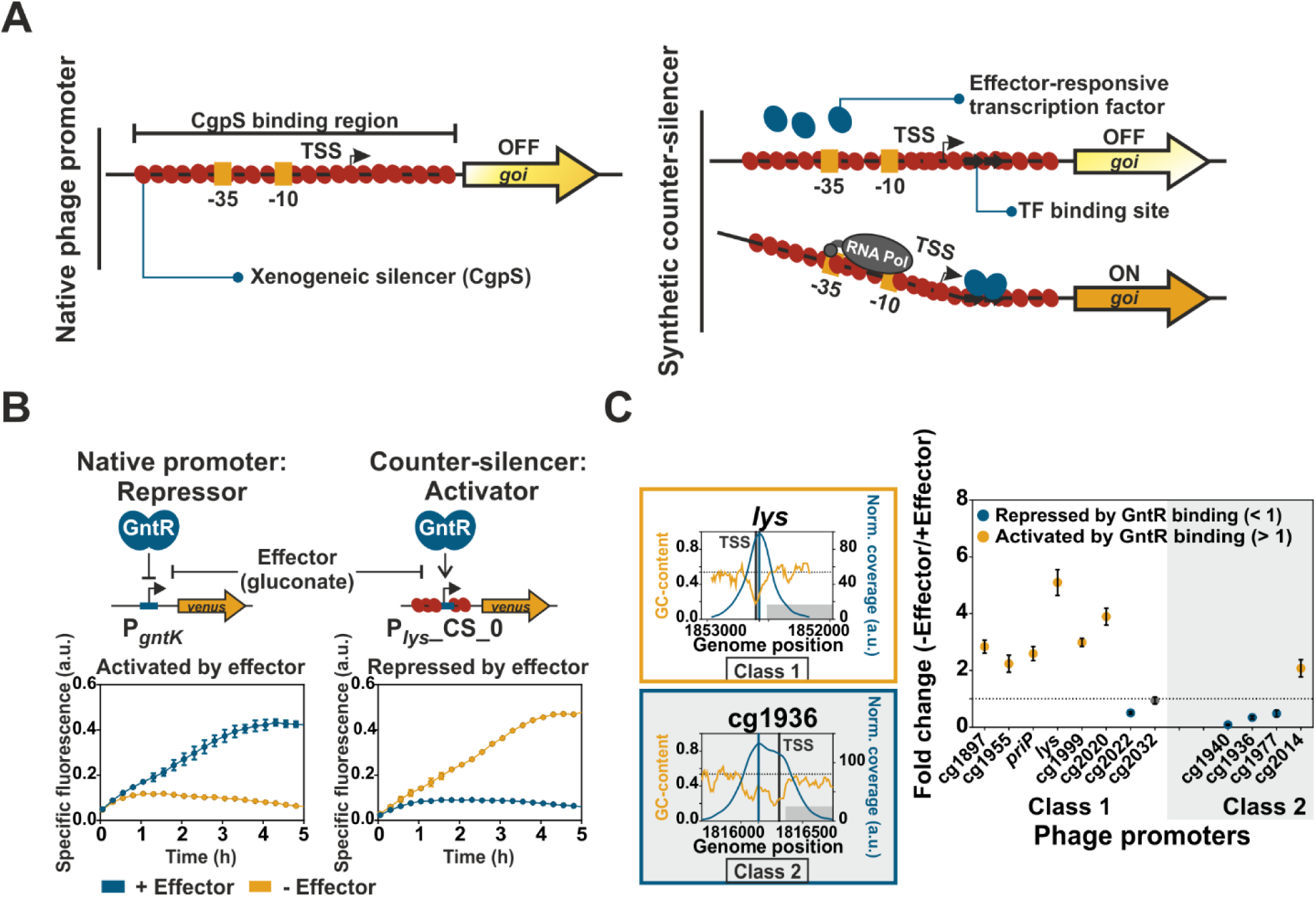
A synthetic approach to study disruptive counter-silencing. **A)** Schematic overview of a native CgpS target promoter (phage) and the corresponding synthetic counter-silencer construct. **B)** Signal inversion by synthetic counter-silencing. Comparison of the reporter outputs of P*_gntK_*, the native target promoter of the regulator of gluconate catabolism GntR (43), and the synthetic GntR-dependent counter-silencer promoter P*_lys_*_CS_0. *C. glutamicum* wild type strains harboring the plasmid-based constructs (pJC1) were cultivated in the absence of the effector (111 mM glucose) or in its presence (100 mM gluconate) in a microtiter cultivation system. Graphs represent the means and error bars the standard deviations of biological triplicates. Backscatter and fluorescence were measured at 15 min intervals. **C)** Counter-silencing efficiency of different phage promoters with inserted GntR binding sites located directly upstream of the position of maximal CgpS binding. Promoters were grouped into two classes based on the width of the region bound by CgpS (class 1 promoters: 500-850 bp, often one distinct drop in GC-profile; class 2 promoters: >850 bp, often broader or multiple drops in GC-content). CgpS coverage and GC-profiles of two representative promoters are shown. Highest ranked TSS are marked as vertical grey lines and position of maximal CgpS binding as vertical blue lines. GC-profiles of all used phage promoters are shown in Figure 1D. *C. glutamicum* wild type cells harboring the plasmid-based counter silencers were cultivated in the presence (100 mM gluconate) or absence (100 mM glucose) of gluconate in a microtiter cultivation system. Fold change ratio of reporter outputs in the absence and in the presence of the effector were calculated based on the specific reporter outputs after five hours of cultivation (Figure S1A). Dots represent the means and error bars the standard deviations of at least biological triplicates. Yellow dots demonstrate counter-silencing (activated by GntR binding), while blue dots represent repression (repressed by GntR binding). Promoters, which did not show significant changes in reporter output, are shown as grey dots (t-test: p-value < 0.05).

Monitoring of fluorescent outputs driven by phage-based synthetic promoter constructs allows the in vivo analysis of silencing and counter-silencing efficiencies. GntR operator sites were indeed confirmed as suitable candidates for the construction of counter-silencers, since the insertion into different phage promotors led to only slightly increased background expression levels in the wild type strain in the presence of gluconate (Figure S1A). The insertion of a GntR binding site (BS) within the CgpS silenced phage promoter P*_lys_* (P*_lys_*_CS_0) led to effector-dependent reporter outputs. GntR binding resulted in an increased reporter output of the counter-silencer construct P*_lys_*_CS_0 when glucose was added as carbon source, while gluconate triggered the dissociation of GntR leading to silencing of promoter activity by CgpS (Figure 3B). This is especially remarkable when considering that the binding site was inserted at the position of maximal CgpS coverage close to the annotated TSS (27 bp downstream (Table S1)). Based on textbook knowledge, this position would rather fit to a repressor function (44, 45). In the case of P*_gntK_*, the GntR binding site overlaps with the TSS leading to repression of gene expression (43). In the context of xenogeneic silencing, however, GntR binding appeared to efficiently interfere with CgpS silencing. Thus, in contrast to the native GntR target P*_gntK_*, the synthetic P*_lys_*-counter-silencer promoter was activated in the absence of gluconate. Although both promoters (P*_gntK_* and P*_lys_*_CS_0) were completely different and had only the 15 bp long GntR binding site in common, they showed very similar, but inverted response to gluconate availability (Figure 3B). This demonstrates the potential of the counter-silencing principle to convert a repressor to an activating, tunable counter-silencer, thereby facilitating the expansion of regulatory networks.

### Disruptive counter-silencing is most efficient at the CgpS nucleation site

To systematically assess the constraints of counter-silencing, 12 phage promoters (eight class 1 and four class 2) were selected as targets to test the efficiency of synthetic counter-silencing. The GntR binding site was inserted directly upstream of the previously identified position of maximal CgpS binding obtained from ChAP-seq analysis (4). To study counter-silencing efficiency, all constructs were analyzed in *C. glutamicum* wild type cells in the presence and absence of the effector molecule gluconate. The ratio of maximal (-gluconate, GntR binding) and minimal (+gluconate; GntR dissociation) reporter outputs was used to compare the counter-silencing efficiency of the different constructs (Figure 3C, Figure S1). Overall, counter-silencing appeared to be more efficient in class 1 promoters typically featuring a bell shaped CgpS peak and a distinct drop in GC-profile. Here, six out of eight constructs showed an effector-responsive counter-silencing behavior. In contrast, only one of four class 2 promoters was activated by GntR binding. The broader regions bound by CgpS are probably stabilizing the silencer-DNA complex allowing to compensate for the local interference effects caused by GntR binding. The general functionality of promoter variants (Figure S1A) was confirmed in the strain Δphage, where all variants showed a significant fluorescent signal (Figure S1B). Interestingly, GntR binding to P_cg2014_ led to counter-silencing in the wild type but to repression in the Δphage strain suggesting that only the destructive interference between CgpS and GntR facilitates efficient transcription of the downstream gene.

### Silencing is mediated by CgpS binding and counter-silencing depends on GntR binding

CgpS as silencer and GntR as counter-silencer are the two key components of the synthetic counter-silencer approach presented in this study. To confirm their presumed functions, mutant analysis and in vitro binding assays with both proteins were performed. The reporter outputs of the native phage promoter P*_lys_* as well as of the corresponding counter-silencer construct (P*_lys_*_CS_0) were analyzed in *C. glutamicum* wild type cells and different mutant strains. In the wild type, the counter-silencer construct showed the expected increase of reporter output upon GntR binding (-gluconate). In line with the assumed counter-silencing function of GntR, both constructs featured a low reporter output in the strain Δ*gntR* lacking both functionally redundant GntR1 and GntR2 regulators (Figure 4A). To confirm the relevance of the inserted GntR operator sequence, different mutated variants were tested as well. Here, neither the insertion of a randomized ‘operator’ sequence, identical in length and nucleotide composition, nor a mutated operator site, where only one conserved base in the GntR motif was exchanged, led to counter-silencing at the P*_lys_* promoter (Figure S2). These results confirmed that counter-silencing directly depends on GntR binding. However, the insertion of a reverse oriented GntR binding site within the silenced promoter allowed counter-silencing showing that this mechanism does not depend on the directionality of the binding site (Figure S3). P*_lys_* and the corresponding counter-silencer construct showed strongly increased promoter activities in the Δphage strain in the absence of CgpS suggesting that CgpS is responsible for silencing. Gluconate-dependent activating effects on reporter outputs were abolished in the absence of CgpS, indicating that GntR act as counter-silencer rather than as classical activator. Re-integration of the *cgpS* gene into the Δphage strain resulting in Δphage::P*_cgpS_*-*cgpS*, confirmed CgpS as the only factor being responsible for silencing of the native phage promoter P*_lys_* and, thus, emphasized that CgpS function does not depend on further phage-encoded accessory proteins (Figure 4A).

**Figure 4:**
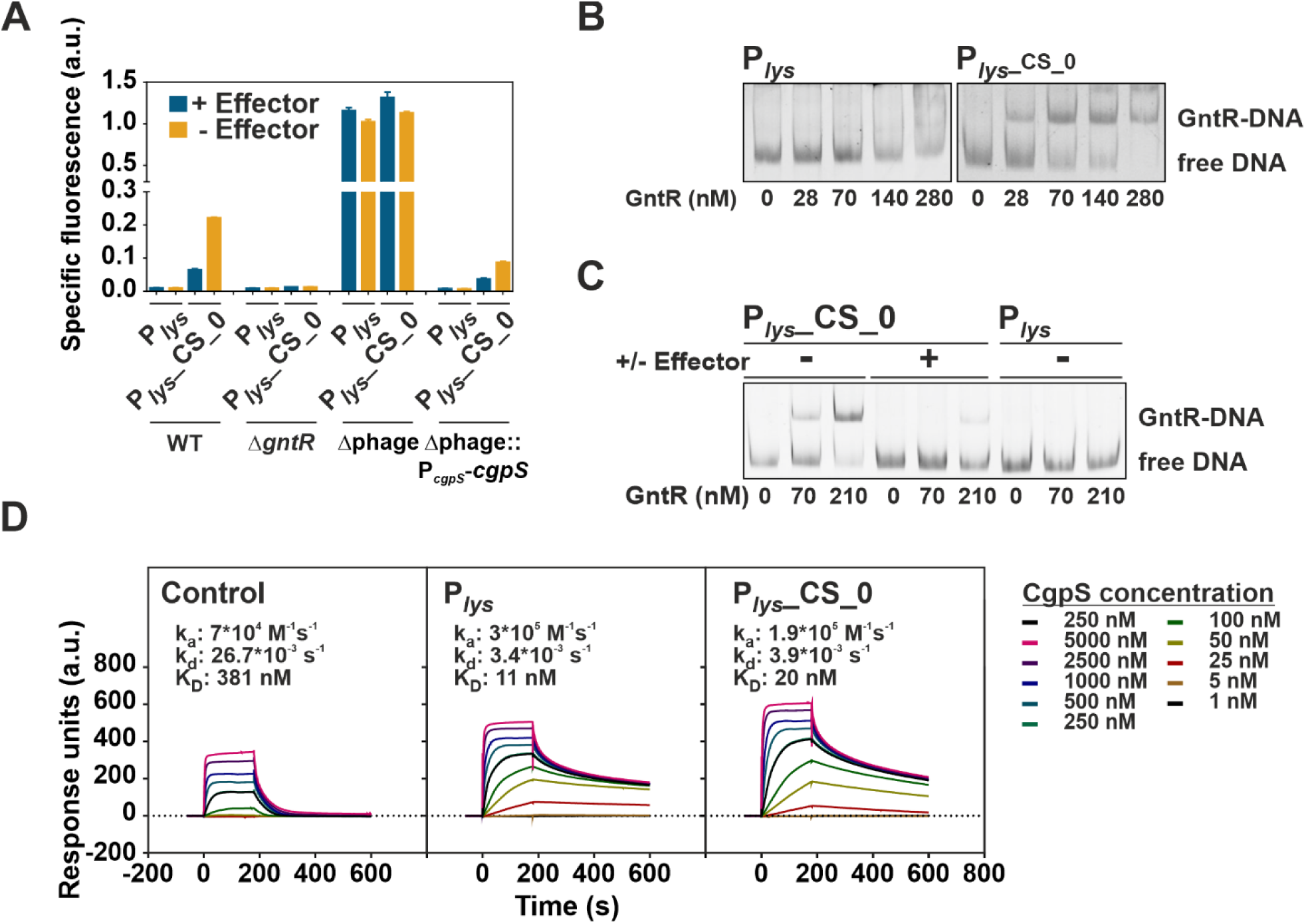
Silencing is mediated by CgpS while counter-silencing depends of GntR binding. **A)** Reporter output (*venus* expression) of different *C. glutamicum* strains carrying the native P*_lys_* promoter or the counter-silencing design P*_lys_*_CS_0 after five hours of cultivation. Both constructs were analyzed in *C. glutamicum* wild type cells, in a *gntR1/2* double deletion strain, in the prophage free strain Δphage (lacking the phage-encoded *cgpS*) and in its variant with re-integrated *cgpS* under control of its native promoter (Δphage::P*_cgpS_*-*cgpS*). Cells were cultivated in a microtiter cultivation system in CGXII medium supplemented with either 100 mM gluconate (+ effector) or 100 mM fructose (-effector). **B)** EMSA analysis of GntR binding to DNA fragments covering the synthetic counter-silencer promoter P*_lys_*_CS_0 (533 bp, 14 nM) or the native phage promoter P*_lys_* (518 bp, 14 nM). **C)** Impact of the effector molecule gluconate on binding of GntR to the synthetic counter-silencer construct. EMSA was performed as described in (B), but GntR and the DNA fragments were incubated either in the presence of the effector (100 mM gluconate) or in its absence (100 mM glucose). **D)** Surface plasmon resonance analysis of CgpS binding kinetics to biotinylated DNA fragments covering the negative control P_cg3336_ (424 bp), the native phage promoter P*_lys_* (424 bp) or the corresponding synthetic counter-silencer construct (439 bp), respectively, that were captured onto a streptavidin-coated sensor-chip. Different concentrations of CgpS were passed over the chip using a contact (association) time of 180 sec, followed by a 420-sec dissociation phase. The increase in RU correlates with an increasing CgpS concentrations.

As a further piece of evidence, electromobility shift assays (EMSA) were performed to confirm the specific binding of GntR to the synthetic counter-silencing construct (P*_lys_*_CS_0) in vitro. In contrast to the native phage promoter, the P*_lys_* fragment containing the GntR operator site showed a significant shift at low GntR concentrations, thus confirming specific GntR binding to P*_lys_*_CS_0 (Figure 4B). Addition of gluconate led to dissociation of GntR (Figure 4C), which is in agreement with previous reports (43). Surface plasmon resonance analysis of CgpS binding to DNA fragments covering either P*_lys_* or the synthetic counter-silencer construct P*_lys_*_CS_0 showed comparable high affinity binding of CgpS to both promoters (K_D_: P*_lys_*: 11 nM, P*_lys_*_CS_0: 20 nM, Figure 4D).

### Impact of operator site position

When analyzing the promoter architecture of horizontally acquired gene clusters, previous studies revealed a certain variability (33). To systematically assess the potential and constraints of the counter-silencing mechanism for evolutionary network expansion, we analyzed the impact of operator site position on counter-silencing efficiency. Therefore, the GntR binding site was inserted at different positions using the prophage promoter P*_lys_* as a test case (Figure 5A). Position 0 is defined as the position located directly upstream of the nucleotide featuring maximal CgpS binding in ChAP-seq studies (4). The position of maximal CgpS binding was located 27 bp downstream of the TSS. *C. glutamicum* wild type cells harboring the plasmid-based constructs (pJC1) were cultivated in the presence or absence of the effector molecule gluconate. Induced and non-induced reporter outputs were strongly influenced by the binding site position. This demonstrated that the inserted binding site itself, depending on its position, already interferes with the silencer-DNA complex (Figure 5B). Comparison of the fold change ratio of reporter outputs in the absence and presence of the effector gluconate revealed that the construct with the GntR binding site located directly upstream of the maximal CgpS binding peak (position 0), showed the highest dynamic range (∼5-fold). This dynamic range decreased when the operator was inserted between 15 bp upstream (-15) and 10 bp downstream (+10) of the maximal CgpS binding peak but constructs still showed counter-silencing in the absence of gluconate (Figure 5C). However, analyzing the -15 promoter variant in the absence of CgpS (Δphage) revealed repression caused by GntR binding demonstrating again that the observed counter-silencing effect is a result of regulatory interference (Figure S4). GntR binding sites located at greater distances led in most cases to relatively low reporter outputs. Here, the expression level tended to be higher when GntR binding was inhibited suggesting GntR acting mainly as a repressor of gene expression at these positions (Figure 5B, C). A similar trend was observed for the phage promoters P_cg1999_ (Figure S5). Altogether, these results demonstrated that the impact of GntR binding on promoter activity strongly depends on the context of xenogeneic silencing. While interference with CgpS binding triggered promoter activation by counter-silencing, GntR binding in the absence of CgpS often lowered the reporter output. Analysis of reporter outputs driven by 5’-truncated promoter variants of P*_lys_* and P*_lys_*_CS_0 revealed that the region >89 bp upstream of the maximal CgpS binding peak is not required for silencing or counter-silencing (Figure S6).

**Figure 5:**
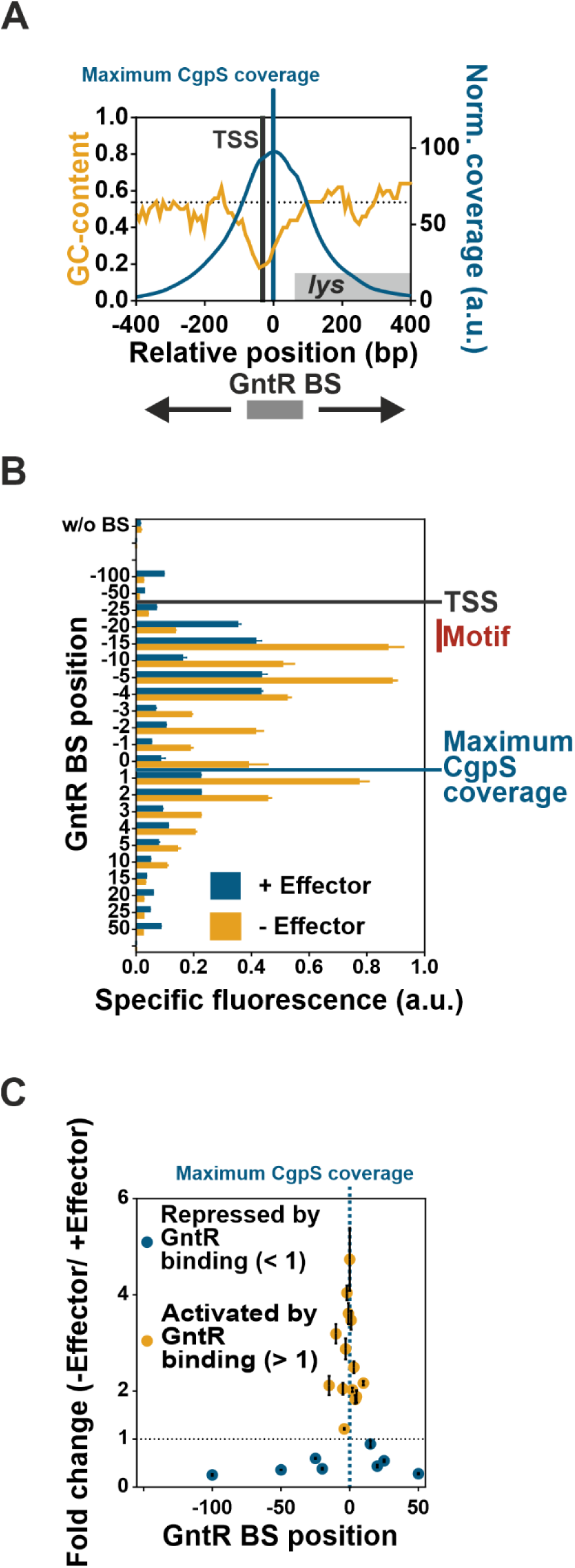
Impact of GntR operator position on inducibility of P*_lys_*-based promoter constructs. **A)** Divergent correlation of GC-profile and CgpS binding coverage (4) of the phage promoter P*_lys_*. The transcriptional start site (TSS) and the position of maximal CgpS binding are shown as vertical lines. Binding site positions (also used in B/C) refer to the sequence base associated with maximal CgpS binding. The position directly upstream of this nucleotide was defined as position 0. **B)** Impact of inserted GntR binding site position on specific reporter outputs in the presence (gluconate) and absence (glucose) of the effector molecule after five hours of cultivation. Positions of TSS and maximal CgpS coverage are marked by horizontal lines and the range of the putative binding motif is shown. **C)** Impact of GntR binding site position on counter-silencing efficiency of P*_lys_*-based promoter constructs. Ratio of specific reporter outputs, shown in (B), were used for the calculation of their inducibility (fold change). Cells harboring the plasmid-based synthetic promoter constructs were grown in CGXII medium supplemented with either 111 mM glucose or 100 mM gluconate. Bars (B) and dots (C) represent the means and error bars the standard deviation of at least biological triplicates.

### Implementation in a genetic toggle switch

The P*_lys_* counter-silencing construct (P*_lys_*_CS_0) and the native GntR target promoter P*_gntK_* showed a very similar promoter output, but an inverted response to gluconate availability. While P*gntK* is repressed by binding of GntR, the counter-silencer promoter is activated by GntR binding in the absence of gluconateB (Figure 3B). Both promoters were combined in a gluconate-dependent, GntR-controlled genetic toggle switch. To monitor the switching between different expression states, the P*_lys_*_CS_0 was fused to the reporter gene *venus*, while the native GntR target promoter P*_gntK_* was fused to the reporter gene *e2-crimson* (Figure 6A). Since the toggle output is only regulated by GntR binding, native GntR levels could be used for toggle control avoiding a negative impact of artificial TF levels on cellular growth. *C. glutamicum* wild type cells harboring the plasmid-based toggle (pJC1) were cultivated in a microfluidic chip device (46) in minimal medium containing either gluconate or glucose as carbon source. The carbon sources were switched after the first 17 hours.

**Figure 6:**
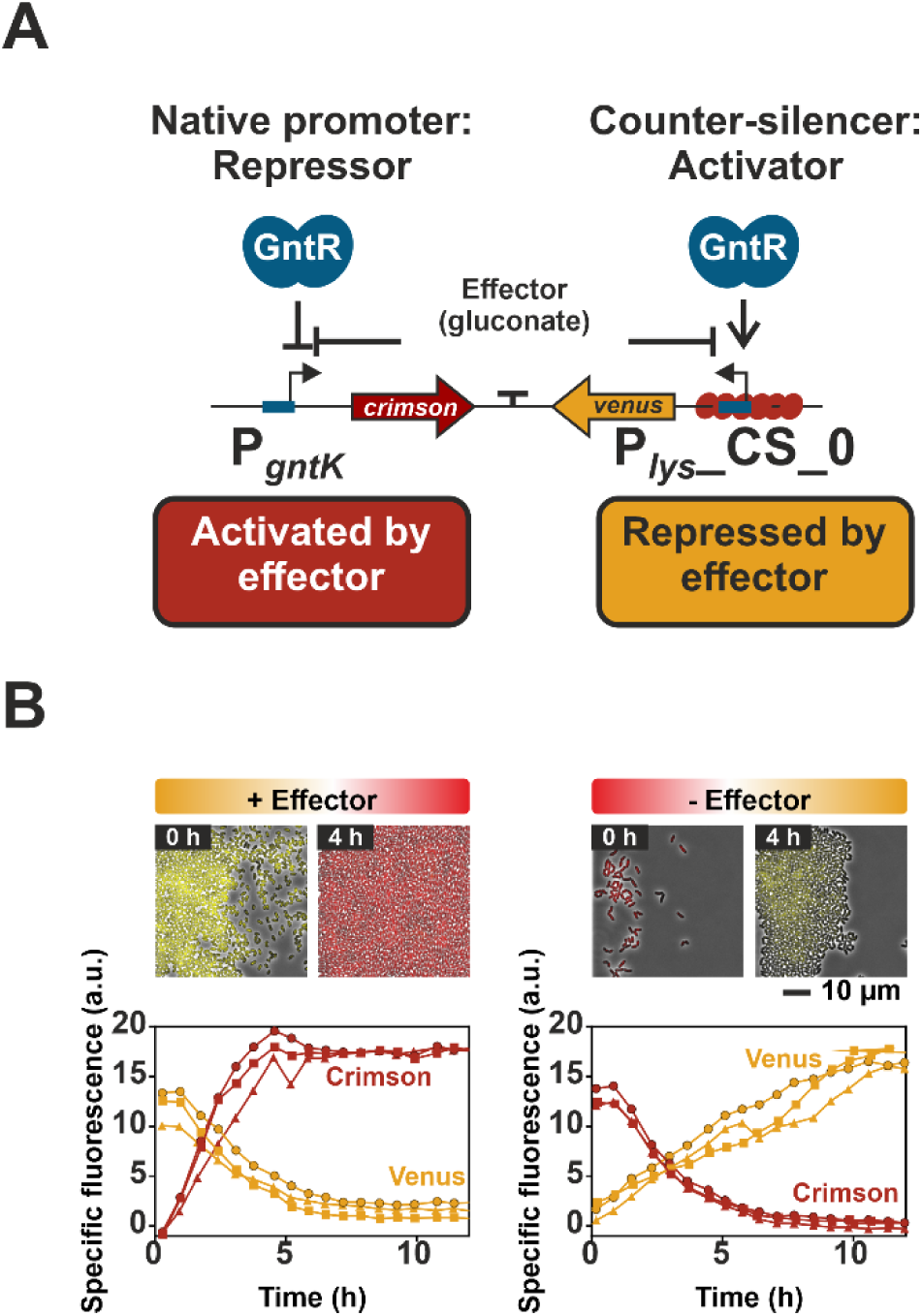
Implementation of P*_gntK_* and P*_lys_*_CS_0 in a genetic toggle switch. **A)** Scheme of the designed toggle switch based on the native GntR target promoter P*_gntK_* and the synthetic GntR-dependent P*_lys_* counter-silencer construct. In order to monitor their activities, the promoters were fused to different reporter genes (P*_lys_*_CS_0-*venus* and P*_gntK_-e2-crimson*). The promoter reporter gene fusions were oriented in opposite directions and separated by a short terminator sequence. **B)** Dynamic switch between both reporter outputs. *C. glutamicum* wild type cells harboring the plasmid-based toggle were cultivated in a microfluidic cultivation system (46) with continuous supply of CGXII medium supplemented either 111 mM glucose or 100 mM gluconate and analyzed by time-lapse microscopy at 20 min intervals. Switch of carbon source supply was performed after first 17 h. This time point was defined as T_0_. The graphs show the specific fluorescence of three independent microcolonies (circles, squares, triangles) over time and images display one representative colony, respectively.

The following time-lapse microscopy analysis revealed that the output of this synthetic toggle is reversible, as shown by the rapid changes in reporter outputs (Figure 6B). This overall design principle allows the control of the toggle by only one specific effector-responsive TF and features a robust and reversible response to effector availability highlighting the potential of this toggle for biotechnological applications, for example, to switch between biomass production and product formation.

## DISCUSSION

The nucleoid-associated protein Lsr2 is conserved throughout the phylum of actinobacteria where it plays an important role in the xenogeneic silencing of horizontally acquired genomic regions (4, 18, 19, 39). The Lsr2-like protein CgpS was recently described as a classical XS protein silencing the expression of cryptic prophage elements and further horizontally acquired elements in *C. glutamicum* (4). In this study, we systematically assessed the promoter architecture of CgpS targets as well as the constraints underlying silencing and counter-silencing of target gene expression. The genome-wide analysis of CgpS-bound regions obtained from ChAP-seq analysis revealed that CgpS targets share a distinct drop in GC-content (Figure 1). The binding to AT-rich DNA is a common feature of XS proteins (3) and was shown to represent an important fitness trait to avoid spurious transcription and the sequestering of the RNA polymerase (8). In vitro protein binding microarray experiments revealed a clear preference of the xenogeneic silencers H-NS, MvaT and Rok for DNA stretches containing flexible TpA steps, while no positive effect of TpA steps was observed for Lsr2 from *Mycobacterium tuberculosis* (20, 47). Here, our ‘design-test-build’ approach, where different synthetic promoter variants were tested in vivo with regard to CgpS-mediated silencing, however, revealed a certain degree of sequence specificity of CgpS towards a binding motif containing A/T steps. While the GC-content was kept constant, alteration of the proposed motif in the P*_lys_* promoter significantly affected in vivo silencing (Figure 2). A scenario which has been proposed for H-NS features a high affinity towards DNA (K_D_: 60 nM) allowing the scanning of DNA until it reaches high affinity sites triggering the nucleation of the tight nucleoprotein complex required for silencing (3, 48, 49). It is important to note, that we almost exclusively relied on in vivo approaches including ChAP-seq analysis and reporter assays to define the parameters affecting silencing and counter-silencing at the systems level. Although in vitro analysis of protein-DNA fragments (often linear DNA) is frequently applied and has provided valuable insights into the binding behavior of XS proteins (20, 47, 49), the conditions do not reflect physiologically relevant situations (DNA topology, protein-protein interaction and interference) and consequently the results have to be interpreted with caution.

While xenogeneic silencing neutralizes the potentially negative effect of invading foreign DNA, counter-silencing allows the integration into the host regulatory network and, thereby, provides access to horizontally acquired genes. This principle has been almost exclusively studied for H-NS in proteobacteria and several types of TFs were shown to counteract H-NS silencing in vivo (15, 50–53). In the case of Lsr2, the mycobacterial iron dependent regulator IdeR represents, to the best of our knowledge, the only example of an investigated Lsr2 counter-silencer. IdeR enables the iron-dependent activation of ferritin by alleviating Lsr2 repression at the *bfrB* locus (54).

During bacterial evolution, mutations leading to the formation of TF operator sequences within silenced promoters allow the controlled expression of the beforehand silenced genes by TF-mediated counter-silencing. In this work, the artificial insertion of the 15 bp short operator sequences of the gluconate-dependent TF GntR within different CgpS target promoters allowed us to study the potential and constraints of counter-silencing of this Lsr2-like XS protein. Binding of GntR to several CgpS target promoters led to transcription initiation of the silenced phage promoters demonstrating that small changes in the DNA sequence added an additional regulatory layer for expression control. Previous studies by Will et al. revealed that counter-silencing and classical activation are different mechanisms of gene regulation (33). While TFs acting as activators typically bind to conserved promoter architectures and promote transcription either by changing the DNA conformation or by recruiting the RNA-polymerase (55), the principle of counter-silencing allows a higher degree of flexibility in terms of promoter architecture (33). For the PhoPQ regulon of *Salmonella enterica* Typhimurium it was shown that PhoP activates core promoters featuring a precise operator position overlapping the -35 box. In contrast, horizontally acquired PhoP target genes show rather diverse promoter architectures and here, transcriptional activation is achieved by counter-silencing of H-NS. In these reported examples the distances between TSS and the closest PhoP binding site vary only by 34 bp (33, 56). This is in a similar range as previous studies of the H-NS target promoter P*_bgl_* revealing a similar window where the insertion of TF operator sites counteract H-NS silencing (34). In the present study, counter-silencing of the Lsr2-like XS CgpS was most efficient at positions close to the position of maximal CgpS binding in a range of approximately 25 bp (Figure 5, Figure S5). By the systematic analysis of truncated promoter variants and by varying the position of TF operator sites, we clearly defined the region required for efficient in vivo silencing and the window where binding of a specific TF may lead to counter-silencing.

In this study, consistently, all tested CgpS target promoters showed significant reporter output in the absence of CgpS confirming that they promote transcription and that CgpS inhibits promoter activity presumably by hindering open complex formation or by trapping the open complex once it has formed (57). Previous studies already suggested that, without xenogeneic silencing, open complex formation is typically not the rate limiting step at AT-rich promoters of horizontally acquired genes, meaning they are constitutive active (33, 58, 59). In the case of the H-NS target promoter *pagC*, an in vitro approach demonstrated RNA-polymerase binding and open complex formation in the absence of additional factors confirming that this promoter alone is transcriptionally competent (33).

In general, two different mechanisms of counter-silencing are conceivable. In one scenario (disruptive mechanism) the interference of TF and XS protein leads to a local disruption of the XS nucleoprotein complex thereby enabling binding of the RNA-polymerase to the DNA. Another possibility would be that counter-silencing allows for supportive contacts between the RNA-polymerase and the TF itself or more distal DNA regions (supportive mechanism). In this study, binding of GntR to the promoter constructs in the absence of the XS CgpS resulted in reduced reporter outputs, although counter-silencing was observed in the wild type. This result strongly speaks for a disruptive rather than a supportive mechanism. It is, nevertheless, intriguing that in the context of xenogeneic silencing binding of a TF at positions close to the TSS leads to promoter activation, where it would cause a road block of transcription at classical promoters (44, 60). These results demonstrate the potential of the counter-silencing principle to convert a repressor to an activating, tunable counter-silencer, thereby facilitating the integration of horizontally acquired DNA into host regulatory networks. Overall, these data illustrate how interference of TFs is shaping global regulatory networks and that the regulatory impact of TF binding is strongly affected by competition with other DNA-binding proteins.

## MATERIAL AND METHODS

### Bacterial strains and cultivation conditions

All bacterial strains and plasmids used in this project are listed in Table S2A-C. The strain *C. glutamicum* ATCC 13032 (61) was used as wild type strain. For detailed information about general growth conditions, microtiter cultivations used to monitor cell growth and fluorescence (62) and about cultivations in the microfluidic chip device (46, 63), the reader is referred to the supplemental methods.

### Recombinant DNA work

All standard molecular methods such as PCR, DNA restriction and Gibson assembly were performed according to previously described standard protocols (65, 66) or according to manufacturer instructions. All plasmids were constructed by Gibson assembly. Details on plasmid construction are provided in Table S2C. DNA sequencing and synthesis of oligonucleotides used for amplification of DNA fragments (inserts for Gibson assembly (Table S2D), biotinylated DNA fragments for surface plasmon resonance (SPR) spectroscopy (Table S2E) and DNA fragments for electrophoretic mobility shift assay (EMSA) (Table S2F)) and sequencing (Table S2G) as well as synthesis of DNA sequences (Table S2H) were performed by Eurofins Genomics (Ebersberg, Germany). Chromosomal DNA of *C. glutamicum* ATCC 13032 was used as PCR template and was prepared as described previously (67). For detailed information about construction of strain Δphage::P*_cgpS_*-*cgpS* via two step homologous recombination (68) and the design of disruptive counter-silencing constructs please see the supplemental methods.

### Determination of transcriptional start sites (TSS)

The determination of transcriptional start sites (TSS) and data analysis was performed with *C. glutamicum* wild type cells by Vertis Biotechnology AG (Vertis Biotechnology AG, Freising, Germany) using the Cappable-seq method developed by Ettwiller and Schildkraut (69). Prophage induction was triggered by adding mitomycin C (MMC). For detailed information about cultivation conditions, RNA preparation and data analysis please see the supplemental methods. Relevant TSS were assigned to phage promoters used in this study when they were located in the promoter region 500 bp upstream of the start codon and directed in gene orientation (Table S1). Multiple TSS assigned to the same promoter were ranked depending on their enrichment scores.

### Plots of CgpS coverage and GC-profiles

Normalized CgpS coverage values obtained from previous ChAP-seq analysis of Pfeifer and colleagues (4) and GC-content of the reference *C. glutamicum* genome BX927147 (70) were plotted to the corresponding genome positions. Both parameters were calculated by a rolling mean with a window size of 50 bp and a step size of 10 bp using R (http://www.R-project.org) (71). The position of maximal CgpS coverage was centered and the promoter orientation was normalized (start codon of the corresponding gene is located on the right site). Identified TSS positions were added. Ends of graphs are defined by the range of the CgpS binding peaks identified in previous ChAP-seq analysis (4). Plotting was performed either by R (71) or by GraphPad prism 7.00 (GraphPad Software, La Jolla. CA. USA).

### Analyses of AT-rich stretches in CgpS binding regions

The *C. glutamicum* genome (BX927147 (70)) was scanned for AT-rich stretches using a custom python script (submitted to GitHub: https://github.com/afilipch/afp/blob/master/genomic/get_at_stretches.py). For further details please see the supplemental methods.

### Protein purification, surface plasmon resonance (SPR) spectroscopy and electrophoretic mobility shift assay (EMSA)

For information about purification of Strep-tagged CgpS and HIS-tagged GntR and the performed in vitro binding assays (surface plasmon resonance (SPR) spectroscopy and electrophoretic mobility shift assay (EMSA)) please see the supplemental methods.

## DATA AVAILABILITY

The custom python script used for scanning for AT-rich stretches is available in the GitHub repository (https://github.com/afilipch/afp/blob/master/genomic/get_at_stretches.py).

## ACKNOWLEDGEMENT

We would like to thank the European Research Council [ERC Starting Grant, grant number 757563]; the Helmholtz Association [grant number W2/W3-096]; and the Deutsche Forschungsgemeinschaft [SPP 1617 grant number FR2759/2-2] for financial support. Furthermore, we would like to thank Dr. Eugen Pfeifer for excellent discussions, for help with R and for the plasmid pK19mobsacB-1199_1201-P*_cgpS_*-*cgpS*. We thank Iska Steffens for her help with the EMSA experiments and Aël Hardy for critical reading of the manuscript. We are very grateful to Prof. Kirsten Jung for using the Bioanalytics core facility of the LMU Munich for SPR analyses.

## SUPPLEMENTAL MATERIAL

**Supplemental methods:** Supplementary information on methods used in this study

### SUPPLEMENTAL TABLES

**Table S1:** TSS analysis of relevant phage promoters after prophage induction with mitomycin C.

**TABLE S2:** Strains, plasmids, oligonucleotides and DNA sequences

**Table S2A:** Strains used in this study.

**Table S2B:** Plasmids from other studies used in this study.

**Table S2C:** Plasmids constructed in this work.

**Table S2D:** Oligonucleotides used in this study for plasmid constructions.

**Table S2E:** Oligonucleotides used for the amplification of DNA probes for surface plasmon resonance analysis.

**Table S2F:** Oligonucleotides used for electrophoretic mobility shift assays.

**Table S2G:** Oligonucleotides used for sequencing.

**Table S2H:** Ordered DNA sequences.

### SUPPLEMENTAL FIGURE LEGENDS

**Figure S1: Counter-silencing of CgpS target promoters. A)** Specific reporter outputs of native phage promoters and corresponding synthetic constructs which were used for the calculation of fold change ratios shown in Figure 3. **B)** To confirm functionality of synthetic promoter variants (GntR BS), which showed only low reporter outputs in the wild type, respective constructs were analyzed in the prophage-free strain Δphage in the absence of the silencer CgpS. All constructs led to significant reporter outputs confirming synthetic promoter functionality (n≥3 biological triplicates).

**Figure S1: Effects of inserted sequences independent of GntR binding. A)** The native GntR BS located within the P*_gntK_* promoter contains four highly conserved and four weakly conserved nucleotides (J. Frunzke, V. Engels, S. Hasenbein, C. Gätgens, and M. Bott, Mol Microbiol, 67:305-322, 2008, doi: 10.1111/j.1365-2958.2007.06020.x). As controls for sequence insertions, which do not allow for GntR binding, control sequences (control-seq) 1 and 2 were inserted at position 0 within the phage promoter P*_lys_*. Control-seq 1 contained the same nucleotide composition as the GntR BS but the sequence was randomized. Control-seq 2 was similar to the GntR BS, but one conserved cytosine was replaced with guanine. Based on previous studies, this exchange was expected to abolish GntR binding (J. Frunzke, V. Engels, S. Hasenbein, C. Gätgens, and M. Bott, Mol Microbiol, 67:305-322, 2008, doi: 10.1111/j.1365-2958.2007.06020.x). **B)** Specific reporter outputs of the promoter variants after five hours of cultivation. Promoter constructs (plasmid backbone pJC1) were fused to the reporter gene *venus* and analyzed in *C. glutamicum* wild type. Cells were cultivated in CGXII in the presence (100 mM gluconate) or absence (100 mM glucose) of the effector molecule gluconate in a microtiter cultivation system. The results shown in this graph demonstrate that mutation of the GntR operator site abolished GntR binding in vivo and that the observed counter-silencing effect strictly depends on GntR binding.

**Figure S2: Effects of directionality of inserted GntR binding site on counter-silencing efficiency.** The 15 bp long GntR BS was inserted either in a forward (P*_lys_*::GntR BS_fw: P*_lys_*_CS_0) or in a reverse orientation (P*_lys_*::GntR BS_rv) in the P*_lys_* promoter (position 0) to analyze if the directionality of the GntR BS influence counter-silencing. Promoter constructs (plasmid backbone pJC1) were fused to the reporter gene *venus* and analyzed in *C. glutamicum* wild type. Cells were cultivated in CGXII medium in the presence (100 mM gluconate) or absence (100 mM glucose) of the effector molecule gluconate in a microtiter cultivation system. Presented is the mean of the specific fluorescence after five hours of cultivation (n≥3 biological triplicates). These results demonstrated that counter-silencing does not depend on the directionality of GntR operator sites.

**Figure S4: It’s all about context – Impact of GntR binding on the output of the P*_lys_* promoter in the presence or absence (Δphage) of CgpS.** Reporter outputs (*venus* expression) driven by a P*_lys_* promoter with an inserted GntR operator sequences 15 bp upstream of the maximal CgpS binding peak (14 bp downstream of TSS) in *C. glutamicum* wild type cells and in the absence of CgpS in the strain Δphage. Cells were cultivated in CGXII in the presence (100 mM gluconate) or absence (111 mM glucose) of the effector molecule gluconate in a microtiter cultivation system. Shown are mean values and standard deviation of reporter outputs of biological triplicates after five hours of cultivation.

**Figure S3: Impact of GntR binding site (BS) position on inducibility of P_cg1999_-based promoter constructs. A)** Divergent correlation of GC-profile and CgpS binding coverage of the phage promoter P_cg1999_. The highest scored transcriptional start site (TSS) and the position of maximal CgpS binding affinity (E. Pfeifer, M. Hünnefeld, O. Popa, T. Polen, D. Kohlheyer, M. Baumgart, and J. Frunzke, Nucleic Acids Res, 44:10117-10131, 2016, doi: 10.1093/nar/gkw692) are shown as vertical lines. BS positions refer to the sequence base associated with maximal CgpS binding peak. The position directly upstream of this nucleotide was defined as position 0. **B)** Impact of inserted GntR BS position on specific reporter outputs in the presence (gluconate) and absence (glucose) of the effector gluconate. Positions of TSS and maximal CgpS coverage are marked by lines. **C)** Impact of GntR binding site position on counter-silencing efficiency of P_cg1999_-based promoter constructs. Ratio of specific reporter outputs, shown in (B), were used for the calculation of their inducibility (fold change). Cells harboring the plasmid-based synthetic promoter constructs were grown in CGXII medium supplemented with either 100 mM glucose or 100 mM gluconate. Bars (B) and dots (C) represent the means and error bars the standard deviation of biological triplicates.

**Figure S4: Definition of the minimal region required for silencing. A)** Reporter outputs (*venus* expression) driven by 5’-truncated promoter versions (-300 and -350 bp truncations) of P*_lys_* and P*_lys_*_CS_0 were analyzed regarding silencing and counter-silencing efficiency. 5’-ends of full-length constructs coincided with the upstream end of the CgpS binding peak (P*_lys_*: 389 bp, P*_lys_*_CS_0: 404 bp upstream of the maximal CgpS binding peak). The distance between maximal CgpS binding peak and ATG was in all constructs 85 bp. Full length promoters were compared to variants ending 89 bp (5’-300) or 39 bp (5’-350) upstream of the maximal CgpS binding peak in P*_lys_*. **B and A)** Specific reporter outputs of the promoter variants after five hours of cultivation. Promoter constructs were fused to the reporter gene *venus* (plasmid backbone pJC1) and analyzed in *C. glutamicum* wild type cells (**B**) and in the prophage-free strain Δphage (Δ*cgpS*) (**C**). Cells were cultivated in CGXII in the presence (100 mM gluconate) or absence (100 mM glucose) of the effector molecule gluconate in a microtiter cultivation system. Shown are mean values and standard deviation of biological triplicates after five hours of cultivation. The results shown in this graph demonstrate that the region >89 bp upstream of the maximal CgpS binding peak of the P*_lys_* promoter is neither involved in silencing nor in counter-silencing. Further 50 bp truncation strongly reduced promoter activity of both promoters also in the Δphage strain.

## REFERENCES

1. Ochman H, Lawrence JG, Groisman EA. 2000. Lateral gene transfer and the nature of bacterial innovation. Nature 405:299–304.

2. Dorman CJ. 2014. H-NS-like nucleoid-associated proteins, mobile genetic elements and horizontal gene transfer in bacteria. Plasmid 75:1–11.

3. Navarre WW. 2016. The impact of gene silencing on horizontal gene transfer and bacterial evolution. Adv Microb Physiol 69:157–186.

4. Pfeifer E, Hünnefeld M, Popa O, Polen T, Kohlheyer D, Baumgart M, Frunzke J. 2016. Silencing of cryptic prophages in *Corynebacterium glutamicum*. Nucleic Acids Res 44:10117–10131.

5. Vogan AA, Higgs PG. 2011. The advantages and disadvantages of horizontal gene transfer and the emergence of the first species. Biol Direct 6:1.

6. Park C, Zhang J. 2012. High expression hampers horizontal gene transfer. Genome Biol Evol 4:523–532.

7. Baltrus DA. 2013. Exploring the costs of horizontal gene transfer. Trends Ecol Evol 28:489–495.

8. Lamberte LE, Baniulyte G, Singh SS, Stringer AM, Bonocora RP, Stracy M, Kapanidis AN, Wade JT, Grainger DC. 2017. Horizontally acquired AT-rich genes in *Escherichia coli* cause toxicity by sequestering RNA polymerase. Nat Microbiol 2:16249.

9. Doron S, Melamed S, Ofir G, Leavitt A, Lopatina A, Keren M, Amitai G, Sorek R. 2018. Systematic discovery of antiphage defense systems in the microbial pangenome. Science 359:eaar4120.

10. Labrie SJ, Samson JE, Moineau S. 2010. Bacteriophage resistance mechanisms. Nat Rev Microbiol 8:317–327.

11. Sorek R, Lawrence CM, Wiedenheft B. 2013. CRISPR-mediated adaptive immune systems in bacteria and archaea. Annu Rev Biochem 82:237–266.

12. Roberts RJ. 2005. How restriction enzymes became the workhorses of molecular biology. Proc Natl Acad Sci U S A 102:5905–5908.

13. Navarre WW, McClelland M, Libby SJ, Fang FC. 2007. Silencing of xenogeneic DNA by H-NS-facilitation of lateral gene transfer in bacteria by a defense system that recognizes foreign DNA. Genes Dev 21:1456–1471.

14. Navarre WW, Porwollik S, Wang Y, McClelland M, Rosen H, Libby SJ, Fang FC. 2006. Selective silencing of foreign DNA with low GC content by the H-NS protein in *Salmonella*. Science 313:236–238.

15. Heroven AK, Nagel G, Tran HJ, Parr S, Dersch P. 2004. RovA is autoregulated and antagonizes H-NS-mediated silencing of invasin and *rovA* expression in *Yersinia pseudotuberculosis*. Mol Microbiol 53:871–888.

16. Oshima T, Ishikawa S, Kurokawa K, Aiba H, Ogasawara N. 2006. *Escherichia coli* histone-like protein H-NS preferentially binds to horizontally acquired DNA in association with RNA polymerase. DNA Res 13:141–153.

17. Tendeng C, Soutourina OA, Danchin A, Bertin PN. 2003. MvaT proteins in *Pseudomonas spp*.: a novel class of H-NS-like proteins. Microbiology 149:3047–3050.

18. Gordon BR, Imperial R, Wang L, Navarre WW, Liu J. 2008. Lsr2 of *Mycobacterium* represents a novel class of H-NS-like proteins. J Bacteriol 190:7052–7059.

19. Gordon BR, Li Y, Wang L, Sintsova A, van Bakel H, Tian S, Navarre WW, Xia B, Liu J. 2010. Lsr2 is a nucleoid-associated protein that targets AT-rich sequences and virulence genes in *Mycobacterium tuberculosis*. Proc Natl Acad Sci U S A 107:5154–5159.

20. Duan B, Ding P, Hughes TR, Navarre WW, Liu J, Xia B. 2018. How bacterial xenogeneic silencer rok distinguishes foreign from self DNA in its resident genome. Nucleic Acids Res 46:10514–10529.

21. Smits WK, Grossman AD. 2010. The transcriptional regulator Rok binds A+T-rich DNA and is involved in repression of a mobile genetic element in *Bacillus subtilis*. PLoS Genet 6:e1001207.

22. Gordon BR, Li Y, Cote A, Weirauch MT, Ding P, Hughes TR, Navarre WW, Xia B, Liu J. 2011. Structural basis for recognition of AT-rich DNA by unrelated xenogeneic silencing proteins. Proc Natl Acad Sci U S A 108:10690–10695.

23. Castang S, Dove SL. 2010. High-order oligomerization is required for the function of the H-NS family member MvaT in *Pseudomonas aeruginosa*. Mol Microbiol 78:916–931.

24. Daubin V, Lerat E, Perrière G. 2003. The source of laterally transferred genes in bacterial genomes. Genome Biol 4:R57.

25. Hildebrand F, Meyer A, Eyre-Walker A. 2010. Evidence of selection upon genomic GC-content in bacteria. PLoS Genet 6:e1001107.

26. Zamenhof S, Brawerman G, Chargaff E. 1952. On the desoxypentose nucleic acids from several microorganisms. Biochim Biophys Acta 9:402–405.

27. Spurio R, Falconi M, Brandi A, Pon CL, Gualerzi CO. 1997. The oligomeric structure of nucleoid protein H-NS is necessary for recognition of intrinsically curved DNA and for DNA bending. EMBO J 16:1795–1805.

28. Winardhi RS, Fu W, Castang S, Li Y, Dove SL, Yan J. 2012. Higher order oligomerization is required for H-NS family member MvaT to form gene-silencing nucleoprotein filament. Nucleic Acids Res 40:8942–8952.

29. Chen JM, Ren H, Shaw JE, Wang YJ, Li M, Leung AS, Tran V, Berbenetz NM, Kocíncová D, Yip CM, Reyrat JM, Liu J. 2008. Lsr2 of *Mycobacterium tuberculosis* is a DNA-bridging protein. Nucleic Acids Res 36:2123–2135.

30. Lim CJ, Lee SY, Kenney LJ, Yan J. 2012. Nucleoprotein filament formation is the structural basis for bacterial protein H-NS gene silencing. Sci Rep 2:509.

31. Landick R, Wade JT, Grainger DC. 2015. H-NS and RNA polymerase: a love-hate relationship? Curr Opin Microbiol 24:53–59.

32. Will WR, Navarre WW, Fang FC. 2015. Integrated circuits: how transcriptional silencing and counter-silencing facilitate bacterial evolution. Curr Opin Microbiol 23:8–13.

33. Will WR, Bale DH, Reid PJ, Libby SJ, Fang FC. 2014. Evolutionary expansion of a regulatory network by counter-silencing. Nat Commun 5:5270.

34. Caramel A, Schnetz K. 1998. Lac and lambda repressors relieve silencing of the *Escherichia coli bgl* promoter. Activation by alteration of a repressing nucleoprotein complex. J Mol Biol 284:875–883.

35. Kane KA, Dorman CJ. 2011. Rational design of an artificial genetic switch: Co-option of the H-NS-repressed *proU* operon by the VirB virulence master regulator. J Bacteriol 193:5950–5960.

36. Colangeli R, Helb D, Vilchèze C, Hazbón MH, Lee C-G, Safi H, Sayers B, Sardone I, Jones MB, Fleischmann RD, Peterson SN, Jacobs WR, Jr., Alland D. 2007. Transcriptional regulation of multi-drug tolerance and antibiotic-induced responses by the histone-like protein Lsr2 in *M. tuberculosis*. PLoS Pathog 3:e87.

37. Colangeli R, Haq A, Arcus VL, Summers E, Magliozzo RS, McBride A, Mitra AK, Radjainia M, Khajo A, Jacobs WR, Salgame P, Alland D. 2009. The multifunctional histone-like protein Lsr2 protects mycobacteria against reactive oxygen intermediates. Proc Natl Acad Sci U S A 106:4414–4418.

38. Pfeifer E, Hünnefeld M, Popa O, Frunzke J. 2019. Impact of xenogeneic silencing on phage-host interactions. J Mol Biol doi:10.1016/j.jmb.2019.02.011.

39. Gehrke EJ, Zhang X, Pimentel-Elardo SM, Johnson AR, Rees CA, Jones SE, Hindra, Gehrke SS, Turvey S, Boursalie S, Hill JE, Carlson EE, Nodwell JR, Elliot MA. 2019. Silencing cryptic specialized metabolism in *Streptomyces* by the nucleoid-associated protein Lsr2. Elife doi:10.7554/eLife.47691.

40. Pfeifer-Sancar K, Mentz A, Rückert C, Kalinowski J. 2013. Comprehensive analysis of the *Corynebacterium glutamicum* transcriptome using an improved RNAseq technique. BMC Genomics 14:888.

41. Bailey TL, Boden M, Buske FA, Frith M, Grant CE, Clementi L, Ren J, Li WW, Noble WS. 2009. MEME Suite: tools for motif discovery and searching. Nucleic Acids Res 37:W202–W208.

42. Grant CE, Bailey TL, Noble WS. 2011. FIMO: scanning for occurrences of a given motif. Bioinformatics 27:1017–1018.

43. Frunzke J, Engels V, Hasenbein S, Gätgens C, Bott M. 2008. Co-ordinated regulation of gluconate catabolism and glucose uptake in *Corynebacterium glutamicum* by two functionally equivalent transcriptional regulators, GntR1 and GntR2. Mol Microbiol 67:305–322.

44. Rojo F. 1999. Repression of transcription initiation in bacteria. J Bacteriol 181:2987–2991.

45. Rydenfelt M, Garcia HG, Cox RS, 3rd, Phillips R. 2014. The influence of promoter architectures and regulatory motifs on gene expression in *Escherichia coli*. PLoS One 9:e114347.

46. Grünberger A, Probst C, Helfrich S, Nanda A, Stute B, Wiechert W, von Lieres E, Nöh K, Frunzke J, Kohlheyer D. 2015. Spatiotemporal microbial single-cell analysis using a high-throughput microfluidics cultivation platform. Cytometry A 87:1101–1115.

47. Ding P, McFarland KA, Jin S, Tong G, Duan B, Yang A, Hughes TR, Liu J, Dove SL, Navarre WW, Xia B. 2015. A novel AT-rich DNA recognition mechanism for bacterial xenogeneic silencer MvaT. PLoS Pathog 11:e1004967.

48. Lang B, Blot N, Bouffartigues E, Buckle M, Geertz M, Gualerzi CO, Mavathur R, Muskhelishvili G, Pon CL, Rimsky S, Stella S, Babu MM, Travers A. 2007. High-affinity DNA binding sites for H-NS provide a molecular basis for selective silencing within proteobacterial genomes. Nucleic Acids Res 35:6330–6337.

49. Gulvady R, Gao Y, Kenney LJ, Yan J. 2018. A single molecule analysis of H-NS uncouples DNA binding affinity from DNA specificity. Nucleic Acids Res 46:10216–10224.

50. Dillon SC, Espinosa E, Hokamp K, Ussery DW, Casadesús J, Dorman CJ. 2012. LeuO is a global regulator of gene expression in *Salmonella enterica* serovar Typhimurium. Mol Microbiol 85:1072–1089.

51. Yu RR, DiRita VJ. 2002. Regulation of gene expression in *Vibrio cholerae* by ToxT involves both antirepression and RNA polymerase stimulation. Mol Microbiol 43:119–134.

52. Navarre WW, Halsey TA, Walthers D, Frye J, McClelland M, Potter JL, Kenney LJ, Gunn JS, Fang FC, Libby SJ. 2005. Co-regulation of *Salmonella enterica* genes required for virulence and resistance to antimicrobial peptides by SlyA and PhoP/PhoQ. Mol Microbiol 56:492–508.

53. Shimada T, Bridier A, Briandet R, Ishihama A. 2011. Novel roles of LeuO in transcription regulation of *E. coli* genome: antagonistic interplay with the universal silencer H-NS. Mol Microbiol 82:378–397.

54. Kurthkoti K, Tare P, Paitchowdhury R, Gowthami VN, Garcia MJ, Colangeli R, Chatterji D, Nagaraja V, Rodriguez GM. 2015. The mycobacterial iron-dependent regulator IdeR induces ferritin (*bfrB*) by alleviating Lsr2 repression. Mol Microbiol 98:864–877.

55. Lee DJ, Minchin SD, Busby SJ. 2012. Activating transcription in bacteria. Annu Rev Microbiol 66:125–152.

56. Zwir I, Latifi T, Perez JC, Huang H, Groisman EA. 2012. The promoter architectural landscape of the *Salmonella* PhoP regulon. Mol Microbiol 84:463–485.

57. Shin M, Song M, Rhee JH, Hong Y, Kim YJ, Seok YJ, Ha KS, Jung SH, Choy HE. 2005. DNA looping-mediated repression by histone-like protein H-NS: specific requirement of Esigma70 as a cofactor for looping. Genes Dev 19:2388–2398.

58. Jordi BJ, Higgins CF. 2000. The downstream regulatory element of the *proU* operon of *Salmonella typhimurium* inhibits open complex formation by RNA polymerase at a distance. J Biol Chem 275:12123–12128.

59. Nagarajavel V, Madhusudan S, Dole S, Rahmouni AR, Schnetz K. 2007. Repression by binding of H-NS within the transcription unit. J Biol Chem 282:23622–23630.

60. Sanchez A, Osborne ML, Friedman LJ, Kondev J, Gelles J. 2011. Mechanism of transcriptional repression at a bacterial promoter by analysis of single molecules. EMBO J 30:3940–3946.

61. Kinoshita S, Udaka S, Shimono M. 1957. Studies on the amino acid fermentation. Part 1. Production of L-glutamic acid by various microorganisms. J Gen Appl Microbiol 3:193–205.

62. Kensy F, Zang E, Faulhammer C, Tan RK, Büchs J. 2009. Validation of a high-throughput fermentation system based on online monitoring of biomass and fluorescence in continuously shaken microtiter plates. Microb Cell Fact 8:31.

63. Grünberger A, Paczia N, Probst C, Schendzielorz G, Eggeling L, Noack S, Wiechert W, Kohlheyer D. 2012. A disposable picolitre bioreactor for cultivation and investigation of industrially relevant bacteria on the single cell level. Lab Chip 12:2060–2068.

64. Keilhauer C, Eggeling L, Sahm H. 1993. Isoleucine synthesis in *Corynebacterium glutamicum*: molecular analysis of the *ilvB*-*ilvN*-*ilvC* operon. J Bacteriol 175:5595–5603.

65. Sambrook J, Russel WD. 2001. Molecular Cloning: A Laboratory Manual (3rd edition). Cold Spring Harbor Laboratory Press, Cold Spring Harbor, NY.

66. Gibson DG, Young L, Chuang RY, Venter JC, Hutchison CA, 3rd, Smith HO. 2009. Enzymatic assembly of DNA molecules up to several hundred kilobases. Nat Methods 6:343–345.

67. Eikmanns BJ, Thum-Schmitz N, Eggeling L, Lüdtke KU, Sahm H. 1994. Nucleotide sequence, expression and transcriptional analysis of the *Corynebacterium glutamicum gltA* gene encoding citrate synthase. Microbiology 140:1817–1828.

68. Niebisch A, Bott M. 2001. Molecular analysis of the cytochrome *bc*_1_-*aa*_3_ branch of the *Corynebacterium glutamicum* respiratory chain containing an unusual diheme cytochrome *c*_1_. Arch Microbiol 175:282–294.

69. Ettwiller L, Buswell J, Yigit E, Schildkraut I. 2016. A novel enrichment strategy reveals unprecedented number of novel transcription start sites at single base resolution in a model prokaryote and the gut microbiome. BMC Genomics 17:199.

70. Kalinowski J, Bathe B, Bartels D, Bischoff N, Bott M, Burkovski A, Dusch N, Eggeling L, Eikmanns BJ, Gaigalat L, Goesmann A, Hartmann M, Huthmacher K, Krämer R, Linke B, McHardy AC, Meyer F, Möckel B, Pfefferle W, Pühler A, Rey DA, Rückert C, Rupp O, Sahm H, Wendisch VF, Wiegräbe I, Tauch A. 2003. The complete *Corynebacterium glutamicum* ATCC 13032 genome sequence and its impact on the production of L-aspartate-derived amino acids and vitamins. J Biotechnol 104:5–25.

71. R Development Core Team. 2016. R: A language and environment for statistical computing, R Foundation for Statistical Computing, Vienna, Austria.

